# Acute Glucagon-Like Peptide-1 Receptor Agonist Liraglutide Prevents Drug-Induced Heroin Seeking in Rats

**DOI:** 10.1101/2021.09.15.460527

**Authors:** Joaquin E. Douton, Nikhil K. Acharya, Brooke Stoltzfus, Dongxiao Sun, Patricia S. Grigson, Jennifer E. Nyland

## Abstract

Substance use disorder is a difficult disease to treat due to its relapsing nature. In the last decade, opioid use disorder has been a threat to public health, being declared an epidemic by the Centers for Disease Control and Prevention. This is a tragic situation, considering there are currently effective, yet not ideal, treatments to prevent relapse. Recent research has shown that hormones that modulate hunger and satiety also can modulate motivated behavior for drugs of abuse. For example, the short-acting analog of glucagon-like peptide-1 (GLP-1), an incretin hormone that regulates homeostatic feeding, has been shown to reduce responding for rewarding stimuli such as food, cocaine, heroin and nicotine. Here, we tested the acute effects of the long-acting GLP-1 analog, liraglutide, on heroin seeking. We found that, in rats with heroin self-administration experience, subcutaneous (sc) administration of an acute dose of 0.3 mg/kg liraglutide was effective in preventing relapse after exposure to three major precipitators: drug-associated cues, stress, and the drug itself. However, the effects of the drug were contingent upon the pretreatment time, with the drug being fully effective when administered using a 6 h, rather than a 4 h pretreatment time. Finally, we confirmed that the reduction in drug seeking is not due to a locomotor impairment, as liraglutide did not significantly alter performance in a rotarod test. As such, this acute non-opioid treatment may serve as a new and effective bridge to treatment.

## Introduction

The chronic and relapsing nature of substance use disorder (SUD) makes it particularly difficult to treat because relapse can occur even after prolonged periods of abstinence.^1^ It is during this abstinence period that people are more susceptible to overdose.^2^ Accordingly, opioid use disorder (OUD) is a serious threat to public health and has been declared an epidemic by the Centers for Disease Control and Prevention.^3^ This is particularly tragic considering there are effective treatments to help to maintain abstinence and to prevent relapse. Medication assisted treatment (MAT) has been proven to reduce relapse in patients with SUD.^4–6^ That said, current therapeutic options are not ideal. For example, both methadone and buprenorphine carry the stigma of ‘replacing one opioid with another’. Further, the opioid receptor antagonist naloxone requires several days of abstinence prior to use and does not alleviate symptoms of withdrawal, resulting in poor compliance. Most importantly, there currently are no non-opioid treatment options for the alleviation of craving – a major risk factor for relapse. It is therefore imperative to find alternative treatments to assist patients with OUD to maintain abstinence.

The incretin hormone glucagon-like peptide-1 (GLP-1), best known for its regulation of homeostatic feeding, also has been observed to modulate reward-motivated behavior. Specifically, the short acting GLP-1 receptor (GLP-1R) agonist, Exendin-4 (Ex-4), has been shown to reduce responding for normally rewarding stimuli such as food, cocaine, alcohol, nicotine, oxycodone and heroin.^7–14^ Importantly, many GLP-1 analogs already have been approved for the treatment of type-2 diabetes and obesity,^15,16^ and, if found effective in reducing responding for drugs of abuse, can be readily re-purposed to treat alcohol use disorder (AUD) and SUD. Of the numerous formulations of GLP-1R agonists, Ex-4 has been the main focus of study in the context of addiction. Its short half-life,^17^ however, will require patients to receive multiple injections throughout the day for continuous prophylactic protection, what could lead to poor compliance. Less research, though, has focused on the effects of GLP-1 agonist formulations with longer half-lives such as liraglutide or semaglutide on drug seeking behavior. The longer half-life of these compounds makes them better candidates as potential treatments to relieve craving and/or withdrawal and to prevent relapse in OUD. In accordance, we found that daily administration of liraglutide during acquisition of heroin self-administration, abstinence, and test reduced heroin self-administration, reduced escalation of heroin self-administration over trials, and reduced drug-induced reinstatement of heroin seeking behavior in rats^18^. Here, we examine the bioavailability of increasing doses of liraglutide over time and test whether acute treatment with liraglutide can reduce cue-induced heroin seeking and drug-, and stress-induced reinstatement of heroin seeking behavior (i.e., the three main precipitating factors of relapse) in heroin-experienced rats. Thus, should the acute administration of liraglutide prove effective, it may serve as a first non-opioid for the acute treatment of opioid craving and withdrawal, and as such, as a non-opioid bridge to treatment.

## Materials and Methods

The total number of subjects was 84 outbred male Sprague-Dawley rats from Charles River (Wilmington, MA) delivered at approximately 90 days of age, weighing between 300 – 400 g at the beginning of the experiment. All subjects were housed individually in standard, suspended, stainless steel cages. The environment in the animal colony room had controlled temperature (21 °C) and humidity, a 12/12 h light/dark cycle, with the light phase starting at 7:00 am. Following a one-week acclimation period to their home cages, rats were habituated to experimenter handling by daily weighing. Water and food were available ad libitum, except where otherwise noted. All studies were approved by the Pennsylvania State University College of Medicine, Institutional Animal Care and Use Committee and performed in accordance with the National Institutes of Health specifications outlined in the Guide for the Care and Use of Laboratory Animals.

### Jugular and Arterial Catheter Implantation Surgery

Rats were anesthetized with isoflurane (4% induction; 2–3% maintenance) and implanted with either arterial catheters in the carotid artery for repeated blood collection in Experiment 1 (n=3) or intravenous jugular catheters in Experiment 2 (n=45) (Instech Laboratories, Inc., Plymouth Meeting, PA) for drug self-administration as described previously.^19^ Following surgery, rats received subcutaneous (sc) injection of the NSAID, carprofen, as post-operative care for at least two days, and were given a full week to recover. Maintenance of jugular catheter patency included flushing catheters using heparinized saline (0.2 mL of 30 IU/mL heparin) every four days. Catheter patency was verified at the end of each week of drug self-administration and the day before each test day using 0.3 mL of propofol (Diprivan 1%).

### Experiment 1

#### Blood Collection

Three rats implanted with arterial catheters were injected daily with increasing doses of liraglutide. Treatment started with 0.06 mg/kg liraglutide (sc) every 24 hours for three days. On the fourth day, the dose was increased to 0.3 mg/kg for another three days. On the seventh day, the dose was increased to 1.0 mg/kg for another three-day period. Blood (200 μL) was extracted from the arterial catheter prior to liraglutide administration, and then every two hours for a total of 10 hours, on the first day of each new dose. Samples where centrifuged at 1000 x g for 10 minutes at 4 °C, and the plasma (supernatant) was transferred to another tube and analyzed using high performance liquid chromatography and tandem mass spectrometry. For more information about the collection and analysis of blood samples see Supporting Information.

### Experiment 2

#### Self-Administration

##### Apparatus

Experiment 2 was conducted in 24 self-administration chambers (MED Associates, Inc., St. Albans, VT) as previously described.^20^ Each chamber measured 30.5 cm in length, 24.0 cm in width, 29.0 cm in height, and was equipped with two retractable empty sipper tubes that entered the chamber through two holes. A stimulus light was located above each hole, and a lickometer circuit was used to monitor licking on each of two empty spouts. In addition, each chamber was equipped with a house light (25 W), a speaker for white noise and a tone generator. Collection of data and events in the chamber were controlled on-line with a Pentium computer using programs written in the Medstate notation language (MED Associates, Inc., St. Albans, VT).

##### Habituation

Rats (n=45) experienced two days of habituation to the self-administration chambers. On the evening prior to the first habituation session, ad libitum water was removed overnight. The rats then underwent one 5-min habituation session per day for 2 days. Starting 2 hours into the light phase, rats were placed in the self-administration chambers for 5 min. During this 5-min period, water was available in one of the two spouts, starting with the center spout (future ‘inactive’ spout) on the first day and the rightmost spout (future ‘active’ spout) on the second day. In order to maintain proper hydration during habituation, rats received overnight access to 20 mL filtered water at the front of the home cage beginning at 5pm. Ad libitum access to water was resumed after the second habituation session.

##### Heroin Self-administration

Two hours into the light cycle, rats were placed in the self-administration chambers (MED Associates, Inc., St. Albans, VT). At the start of each session, two empty spouts advanced. Contacts with the center spout had no consequence (inactive spout). In contrast, a cue light above the rightmost spout indicated its availability, and completion of a fixed ratio of 10 (FR=10) contacts with this empty spout led to a 6-second intravenous (iv) infusion of either saline (n=20) or 0.06 mg/infusion heroin (n=25). Each infusion was accompanied by a 20-second time-out period in which the cue light turned off, the house light turned on, the empty spouts retracted, and the sound of a tone signaled the time out period. After each timeout period, the active and inactive spouts were again advanced. Rats were allowed the opportunity to self-administer heroin or saline for 6 hours each day for 5 days per week until 11 trials were completed. Once self-administration ended, 7 rats from the heroin condition that did not acquire self-administration behaviors were removed from the experiment.

##### Cue-induced Seeking and Drug-induced Reinstatement Four hours after Liraglutide Treatment

Twenty-four hours after the last self-administration trial, Group 1, consisting of 8 rats from the heroin group and 8 rats from the saline group, were assigned to vehicle (n=8) or liraglutide (n=8) pretreatment conditions. Groups were balanced based on the number of infusions taken during self-administration. At the beginning of the light cycle, rats received a single sc injection of vehicle (saline) or liraglutide (0.3 mg/kg). Four hours later, rats were placed in the self-administration chamber and subjected to a 3-hour extinction test during which time all cues associated with the drug (cue light, tone, etc.) were presented as usual, but contacts with the active spout did not deliver an infusion of saline or drug. Immediately after the third hour, when seeking had extinguished, rats received a single computer-controlled non-contingent iv infusion of saline or heroin (0.06 mg/infusion, depending on their initial group assignment) and reinstatement of heroin seeking behavior was assessed across another hour. The number of contacts with the active spout during the first hour of the three-hour extinction period was interpreted as cue-induced seeking, while the number of contacts during the hour after the single, non-contingent iv infusion was interpreted as drug-induced reinstatement of heroin seeking. Comparison between active and inactive spout contacts was used as a proxy of goal-directed behavior.

##### Cue-induced Seeking, Drug-induced Reinstatement, and Stress-induced Reinstatement Six hours after Liraglutide Pretreatment

The remaining rats (Group 2) received liraglutide (n=10) or vehicle (n=12) pretreatment six hours prior to the extinction/reinstatement session. All other procedures were identical to those described for the four-hour pretreatment group, with the exception of the longer pretreatment time, and an additional test for stress-induced reinstatement. After the first extinction/reinstatement test, these rats were given 14 days of abstinence in the home cage before being subjected to a second extinction/reinstatement test. As with the first test, rats received a sc injection of vehicle (n=12) or liraglutide (0.3 mg/kg, n=10) six hours before the beginning of the test. Thereafter, rats were re-exposed to a 3-hour period of extinction as described. At the end of the third hour, all rats were injected intraperitoneally (ip) with the alpha-2 adrenergic antagonist, yohimbine (0.5 mg/kg), which has been observed to induce anxiety- and stress-like responses in laboratory animals and humans, and to induce reinstatement of drug seeking behavior.^21–23^ Following yohimbine administration, heroin seeking behavior was monitored for two hours.

### Experiment 3

#### Rotarod Locomotor Test

To verify that the effect of liraglutide on behavior during the extinction/reinstatement tests were not due to lethargy or motor impairment, 36 naïve rats underwent a rotarod test to assess the effects of liraglutide on motor coordination and balance. The rotarod test measures the length of time a rat can remain on a rotating rod, and was developed to measure the effects of drugs on motor coordination.^24,25^ The rotarod consists of a rod, 60 mm in diameter, divided into 4 lanes separated by Plexiglas dividers (Panlab, Barcelona, Spain). The speed of rotation and acceleration can be controlled and the floor beneath each lane triggers the timer to stop when a rat falls onto it. In this experiment, rats were habituated to the rotarod apparatus for four days prior to testing. During habituation, rats were placed on the rotarod apparatus with the rod set to rotate at a constant rate of 4 RPM. Each rat was placed on the rotating rod for 5 minutes; rats that fell from the apparatus were placed back onto the rod until the habituation period of 5 minutes was completed. Once the rats were comfortable walking on the rod at 4 RPM, the speed was gradually increased with successive trials until the rats were comfortable walking on the rod at 16 RPM. The final two days of habituation tested the ability of the rats to remain on the rotarod as speed increased from 4 to 40 RPM over a period of 180 seconds. Each rat underwent three of these acceleration trials on each of the two final habituation days. Only rats that consistently achieved a latency of 40 seconds or more on these acceleration trials continued to the liraglutide test (n=14 rats).

On test day, rats received a sc injection of either saline (n=7) or 0.3 mg/kg liraglutide (n=7). The latency for each rat to fall off the accelerating rotarod (4 RPM to 40 RPM over 180 seconds) was assessed at 4, 6, 8, and 10 hours after the injection. This period correlates with the peak concentration of liraglutide as well as the period in which extinction/reinstatement tests were conducted in the previous experiments. The latency to fall off the rotating rod was measured twice at each time point and the greater of the two latencies was selected for each rat. Seventy-two hours later, rats from both groups were assigned to either saline or morphine conditions using a crossover design to balance prior drug experience. Rats then received an ip injection of either saline (n=6) or morphine (15 mg/kg, n=6). The rotarod test was conducted exactly as described for liraglutide, however, due to the short half-life of morphine, the time points tested were 15, 30, 60, and 90 minutes after the injection. Morphine has been shown to reduce locomotor activity at this dose and within this time-frame and was used as a positive control.^26^ Two subjects were removed from the experiment due to lack of cooperation and failure to complete the second test.

### Data Analysis

Differences between groups were analyzed using two-way or mixed factorial Analysis of Variance (ANOVAs) varying group (saline, heroin), pretreatment (vehicle, liraglutide), and trial, or time where applicable. Post hoc analyses were conducted using Tukey’s post hoc tests or Bonferroni correction to account for repeated sampling. Raw latency data were analyzed using the Kaplan-Meier survival curves and the Log-rank (Mantel-Cox) test. All data, including area under the curve (AUC) were analyzed using Prism version 9.1.1., GraphPad Software (La Jolla California USA).

## Results

### Experiment 1

Figure 1 shows the mean plasma concentration of liraglutide over time following the first daily injection of each of the following escalating doses: 0.06, 0.3, and 1.0 mg/kg. Rats were injected daily, and the dose was increased after every third day. Liraglutide plasma concentration, as expected, increased dose-dependently. The starting, or baseline concentration, maximum concentration (Cmax), area under the curve (AUC), and time to peak concentration (Tmax) are listed in Table 1. The area under the curve (AUC_0-10h_) dose dependently increased from 24,437 ngh/mL for the 0.06 mg/kg dose, to 211,575 ngh/mL for the 0.3 mg/kg dose, and 500,686 ngh/mL for the 1.0 mg/kg dose. Importantly, liraglutide was not completely cleared after 24 hours, as the baseline observed for the 0.3 mg/kg and 1.0 mg/kg doses did not return to 0 ng/mL. The first administration of the 0.3 mg/kg dose was given 24 hours after the third daily injection of the 0.06 mg/kg dose, and at the time of the first 0.3 mg/kg dose, the plasma concentration was 4,532 ng/mL; similarly, the liraglutide concentration prior to the first injection of the 1.0 mg/kg dose was 17,160 ng/mL (see Figure 1, 0 hr).

**Figure 1.**
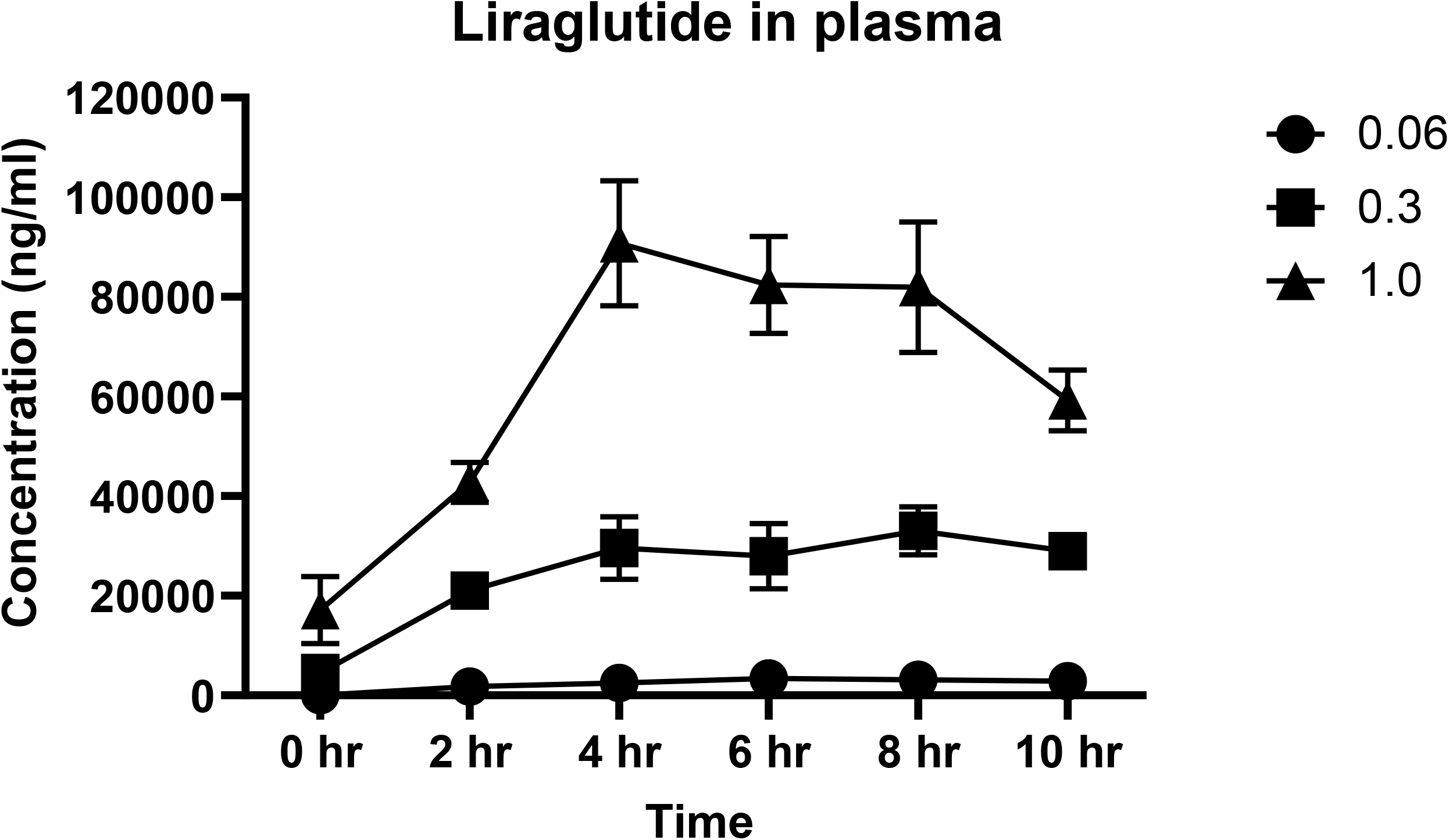
Liraglutide plasma concentration. Rats (n=3) were treated daily with increasing doses of LIR. Blood samples were collected on days 1, 4 and 7, before injection (0h), and at 2, 4, 6, 8 and 10 hours post injection. Figure 1 shows Mean (± SEM) liraglutide concentration (ng/mL) in plasma over ten hours following the first sc injection of 0.06 mg/kg, 0.3 mg/kg, and 1.0 mg/kg doses of liraglutide.

**Table 1.** Pharmacokinetic parameters across doses of liraglutide.

|  | **0.06 mg/kg** | **0.3 mg/kg** | **1.0 mg/kg** |
| --- | --- | --- | --- |
| **Baseline concentration (ng/mL)** | 0 | 4,532 | 17,160 |
| **AUC<sub>0-10</sub> (ngh/mL)** | 24,437 | 211,575 | 500,686 |
| **Tmax (hours)** | 6 | 8 | 4 |
| **Cmax (ng/mL)** | 3,393 | 33,056 | 90,808 |
| **AUC<sub>0-peak</sub> (ngh/mL)** | 11,978 | 158,664 | 124,821 |

AUC: Area Under the Curve; Tmax: Time of maximum concentration; Cmax: Maximum concentration.

### Experiment 2

#### Heroin Self-Administration

Prior to treatment, rats had the opportunity to self-administer either heroin or saline for a period of 6 hours per session, 5 days per week, over 11 trials (see figure 2A). Figure 2B shows the mean number of infusions over the 11 self-administration trials after dividing the rats into future treatment groups of either vehicle or liraglutide for the test day. A 2 × 2 × 11 mixed factorial ANOVA varying group (heroin vs saline), future treatment (vehicle vs liraglutide) and trial (1 – 11) found a significant group x trial interaction, (F_10,340_=12.47, p<0.0001), and post hoc analyses of the interaction indicated that rats in the heroin group took more infusions over time compared with the saline group, and this reached significance for trials 5 through 11 (p<0.05). In addition, rats that self-administered heroin showed clear goal-directed behavior, as the number of contacts with the active spout differed significantly from those observed with the inactive spout, (Figure 2C). This conclusion was supported by post hoc analysis of a significant group by spout interaction of a 2 × 2 × 2 ANOVA varying group, future treatment and spout (active vs inactive - F_1,35_=33.4, p<0.0001).

**Figure 2.**
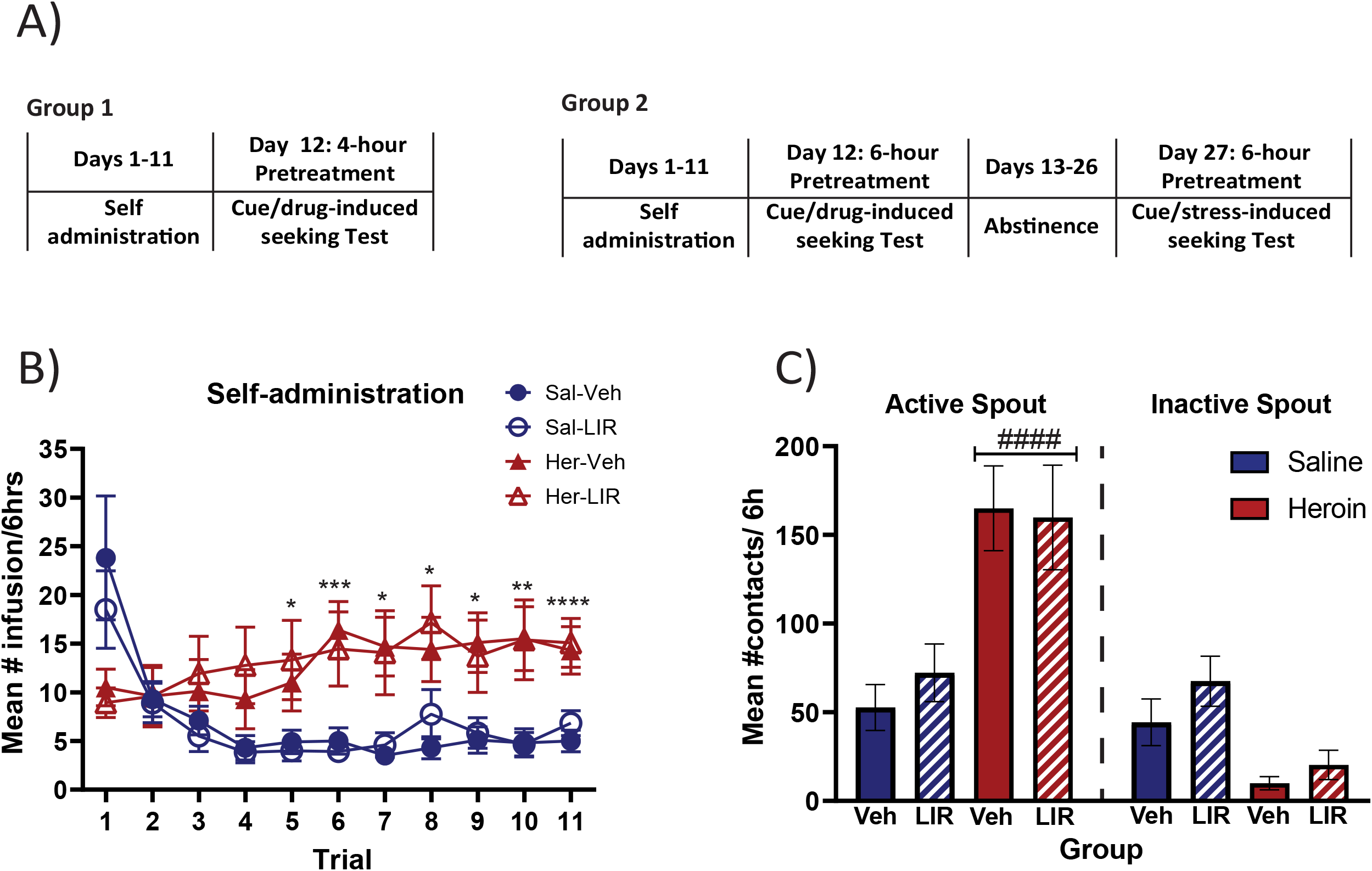
**A.** Timeline of the study. In Group 1, rats had 11 trials of either saline or 0.06mg/infusion heroin self-administration (FR=10). The day after the last self-administration trial, rats were subjected to an extinction reinstatement test in order to assess drug-seeking behavior. Group 2 had the same 11 self-administration trials as test day. However, this group had experienced an additional 14 days of abstinence and a second test that in which cue-induced seeking and stress-induced reinstatement were analyzed. **B. Self-administration.** Mean ± SEM number of infusions/6h across 11 trials for rats that self-administered saline (n=20; blue) or heroin (n=18; red). The groups were also split according to future treatment with either vehicle (close symbols) or 0.03 mg/kg liraglutide (open symbols). **C. Goal-directed behavior.** Mean (± SEM) number of active and inactive spout contacts during the last self-administration trial for rats for rats that self-administered saline (n=20; blue) or heroin (n=18; red). The groups were also split according to future treatment with either vehicle (filled bars) or 0.03 mg/kg liraglutide (hatched symbols). *Significant difference between groups (heroin vs saline). #Significant difference between active and inactive spout. For any given symbol: *p<0.05; **p<0.01; ***p<0.001; ****p<0.0001.

#### Extinction/Reinstatement Test with 4-hour Pretreatment

Four hours after treatment with vehicle or liraglutide (0.3 mg/kg, sc), rats in Group 1 were subjected to 3 hours of extinction followed by a single non-contingent infusion of saline or heroin and one additional hour of drug-induced reinstatement of heroin seeking behavior (see Figure 3A). When treated 4 hours prior to testing (Figure 3B), there was no effect of liraglutide on cue-induced seeking during the first hour of extinction as indicated by the lack of a significant group x treatment interaction (F<1), or main effect of treatment (F_1,12_=4.07, p>0.05) following a 2 × 2 × 2 ANOVA varying group (saline vs heroin), treatment (vehicle vs liraglutide) and spout (active vs inactive). The group x spout interaction, however, was significant (F_1,12_=5.17, p<0.05), showing that rats with heroin experience interacted more with the previously active spout than with the previously inactive spout overall. During the entire extinction session (Figure 3C), a significant group x time interaction of a 2 × 2 × 3 ANOVA (F_2,24_=5.047, p<0.05) showed that heroin-seeking behavior extinguished across the session. Finally, during the drug-induced reinstatement test (Figure 3D), which occurred 7 hours after liraglutide or saline pretreatment, rats in the liraglutide-treated heroin group made significantly fewer contacts with the previously active spout than rats in the vehicle-treated heroin group. In fact, seeking was completely abolished as, akin to rats in the saline groups (both vehicle- and liraglutide-treated), no contacts were observed with the active spout. As a consequence, vehicle-treated rats in the heroin group were the only group that showed a significant difference in contacts between the active and inactive spouts. Results of a 2 × 2 × 2 mixed factorial ANOVA supported these conclusions as indicated by a significant group x treatment x spout interaction (F_1,12_=9.037, p<0.05) and confirmed by post hoc analyses (p<0.05).

**Figure 3.**
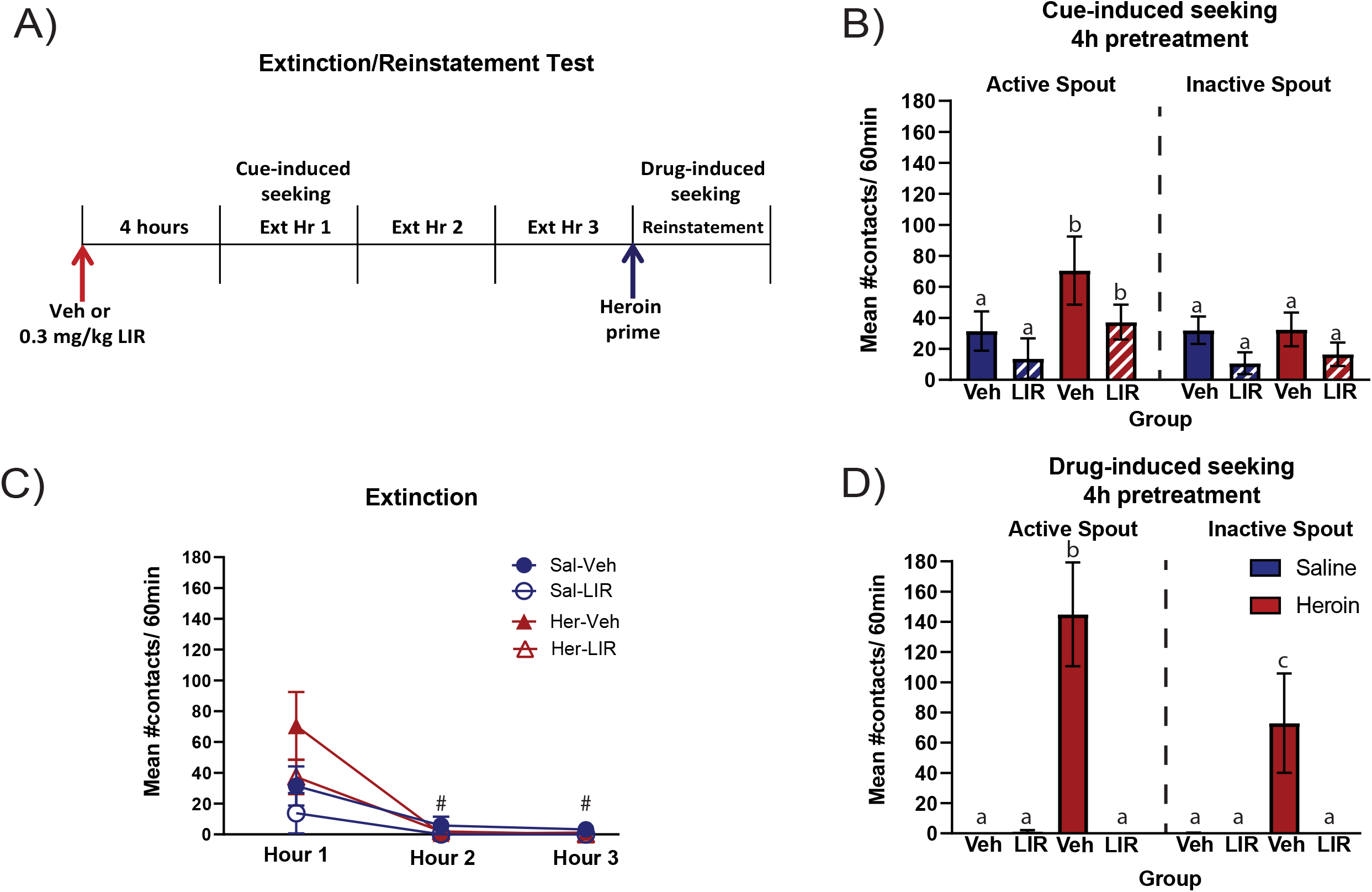
Cue-induced seeking and drug-induced seeking with 4h pretreatment. **A. Extinction/Reinstatement protocol**. Rats were pretreated 4h before the beginning of the test. At test, rats had 3h of extinction in which all the cues associated with the drug were present but contacts with the active spout did not deliver any drug. At the end of hour 3, an iv infusion of saline or 0.06mg/infusion heroin was automatically delivered to reinstate seeking behavior. Cue-induced seeking was evaluated during the first hour of extinction and drug-induced seeking was evaluated during the 1^st^ hour of reinstatement test. **B. Cue-induced seeking.** Mean (± SEM) number of active and inactive spout contacts during the first hour of extinction for rats with a history of saline or heroin self-administration pretreated with vehicle (saline) or (0.3mg/kg) LIR 4 h prior to the start of the test. **C. Extinction.** Mean (± SEM) number of active spout contacts during the three hours of extinction for the same group of rats. **D. Drug-induced seeking.** Mean (± SEM) number of active and inactive spout contacts emitted during reinstatement for the same group of rats. Saline-veh (n=4), saline-LIR (n=4), heroin-veh (n=4), heroin-LIR (n=4). Different letters indicate significant differences. #Significant difference within heroin the group at different times. #p<0.05.

#### Extinction/Reinstatement Test with 6-hour Pretreatment

In contrast to rats that received a 4-hour pretreatment, rats in Group 2, that were pretreated 6 hours prior to the start of the extinction/reinstatement test (see Figure 4A) showed a significant reduction in both cue- and drug-induced seeking (Figure 4B and D). During cue-induced seeking, liraglutide-treated rats in the heroin group made significantly fewer contacts with the previously active spout compared with vehicle-treated rats in the heroin group. No effect of liraglutide was observed in the saline self-administering groups. In addition, vehicle-treated rats in the heroin group were the only ones that showed a significant difference between active and inactive spout contacts. These conclusions were supported by post hoc analysis (p<0.05) of significant group x treatment x spout interaction of a 2 × 2 × 2 ANOVA (F_1,18_=12.1, p<0.01). Moreover, when analyzing the entire extinction period, a significant group x treatment x time interaction (F_2,36_=7.07, p<0.01) showed that vehicle-treated rats with a history of heroin self-administration extinguished seeking behavior by the third hour (Figure 4C). Results of the drug-induced reinstatement test, which occurred 9 hours after treatment with vehicle or liraglutide, were similar to those observed in Group 1 (Figure 4D). Rats with a history of heroin self-administration treated with saline exhibited more contacts with the active spout than the inactive spout and greater drug seeking following the non-contingent iv-heroin infusion, and this behavior was significantly reduced by liraglutide. In support, post hoc analysis of a significant group x treatment x spout interaction (F_1,18_=23.97, p<0.0001) confirmed that, within the heroin group, liraglutide-treated rats made fewer contacts with the active spout than vehicle-treated rats (p<0.05), while no differences were observed in rats with a history of saline self-administration (p>0.05).

**Figure 4.**
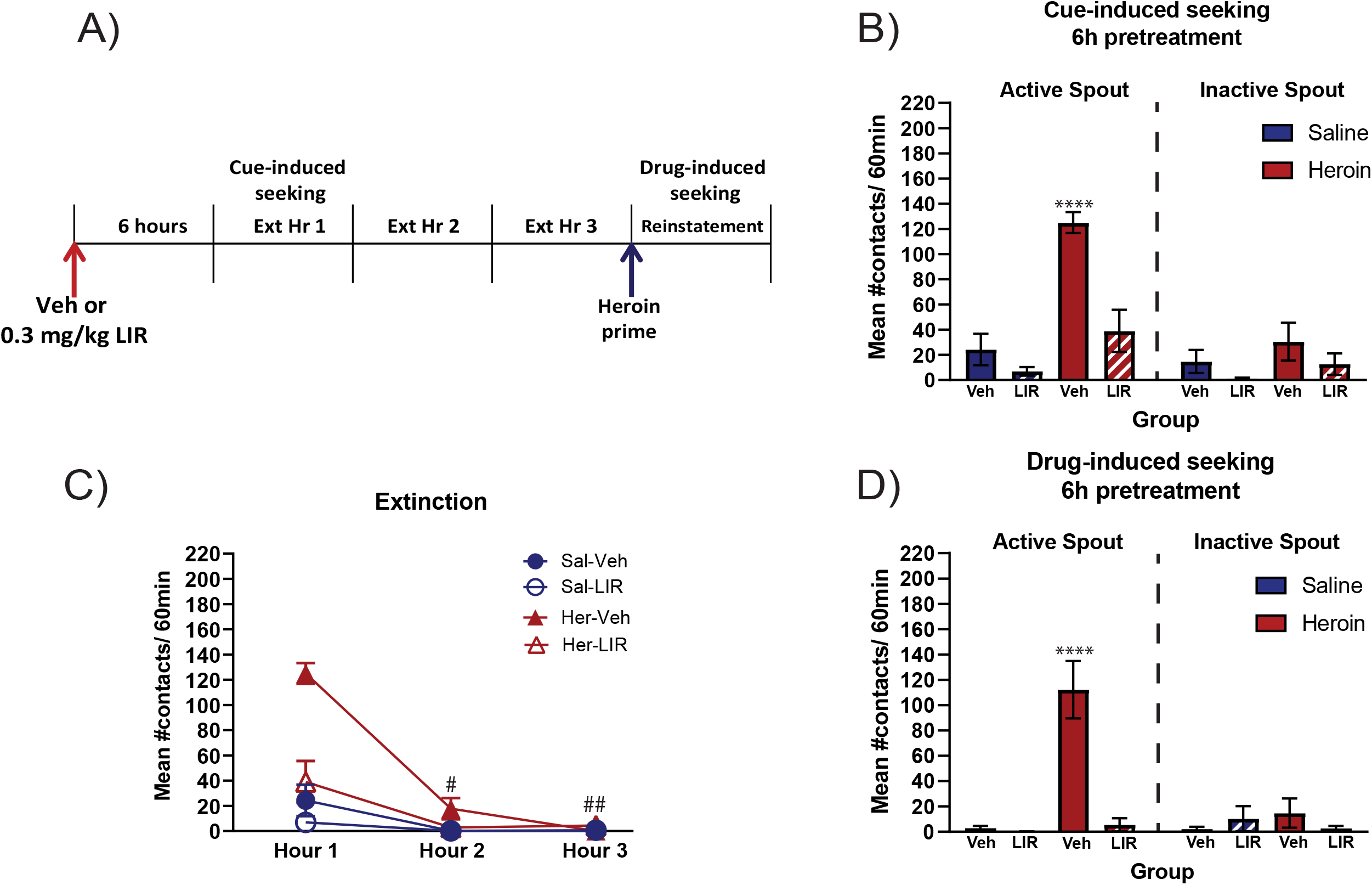
Cue-induced seeking and drug-induced seeking with 6h pretreatment. **A. Extinction/Reinstatement protocol.** Rats were pretreated 6h before the beginning of the test. At test, rats had 3h of extinction in which all the cues associated with the drug were present but contacts with the active spout did not deliver any drug. At the end of hour 3, an iv infusion of saline or 0.06mg/infusion heroin was automatically delivered to reinstate seeking behavior. Cue-induced seeking was evaluated during the first hour of extinction and drug-induced seeking was evaluated during reinstatement. **B. Cue-induced seeking.** Mean (± SEM) number of active and inactive spout contacts during the first hour of extinction for rats with a history of saline or heroin self-administration pretreated with vehicle (saline) or 0.3 mg/kg LIR 6 h prior to the start of the test. **C. Extinction.** Mean (± SEM) number of active spout contacts during the three hours of extinction for the same group of rats. **D. Drug-induced seeking.** Mean (± SEM) number of active and inactive spout contacts emitted during reinstatement for the same group of rats. Saline-veh (n=6), saline-LIR (n=6), heroin-veh (n=4), heroin-LIR (n=6). *Significant differences between her-vehicle active spout contacts and all other conditions. *p<0.05; **p<0.01, ****p<0.0001. #Significant difference within heroin-liraglutide group at different times. #p<0.05; ##p<0.01.

#### Stress-induced seeking

Rats from the 6-hour pretreatment group were given an additional 14-day period of abstinence after the extinction/reinstatement test (see Figure 2A). Following abstinence, cue-induced seeking was re-analyzed along with stress-induced reinstatement (see Figure 5A). Similar to the previous test, a 6-hour pretreatment with 0.3 mg/kg liraglutide reduced cue-induced seeking during the first hour of extinction in rats with history of heroin self-administration (Figure 5B). This was supported by a significant group x treatment x spout interaction (F_1,18_=9.71, p<0.01). Post hoc analyses of the interaction confirmed that the heroin-vehicle group made significantly more contacts on the formerly active spout compared with the other three groups during the first hour of the test (p<0.05). In addition, post hoc analyses (p<0.05) of a significant group x treatment x time interaction across the entire 3 h extinction session (F_2,36_=3.82, p<0.05) confirmed that heroin seeking behavior by the vehicle treated rats with a history of heroin self-administration was extinguished by the last hour of the session (Figure 5C). Finally, 0.3 mg/kg liraglutide also reduced stress-induced reinstatement of heroin seeking behavior (see Figure 5D). Thus, following yohimbine administration, vehicle-treated rats with a history of heroin self-administration showed robust reinstatement of drug-seeking. These effects were eliminated for liraglutide-treated rats with heroin experience. Additionally, only vehicle-treated rats in the heroin group showed a significant difference between active and inactive spouts contacts. These findings were supported by post hoc analyses (p<0.05) of a significant group x treatment x spout interaction (F_1,18_=9.69; p<0.01).

**Figure 5.**
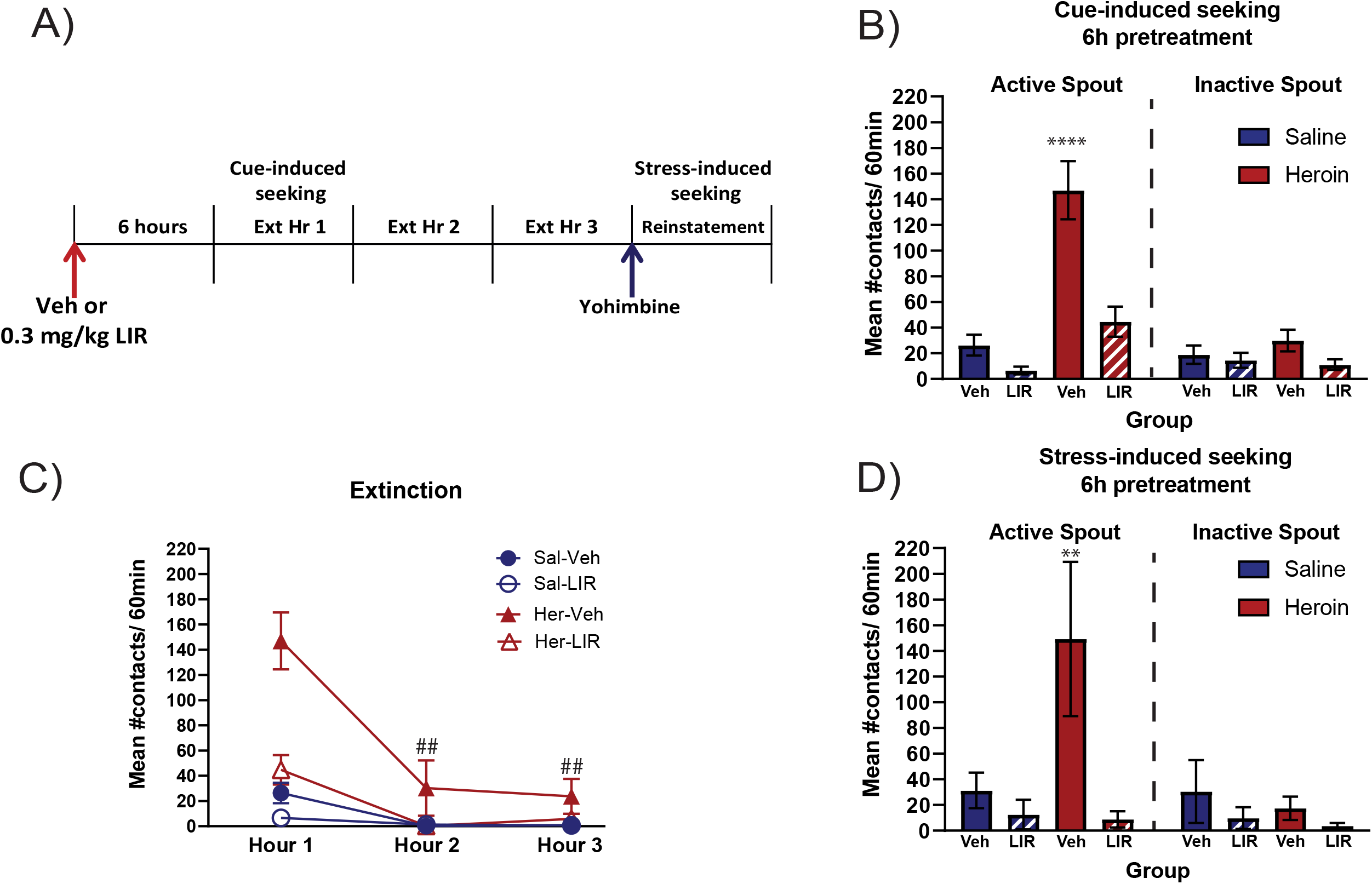
Cue-induced seeking and stress-induced seeking with 6h pretreatment. **A. Extinction/Reinstatement protocol.** Rats were pretreated 6h before the beginning of the test. At test, rats had 3h of extinction in which all the cues associated with the drug were present but contacts with the active spout did not deliver any drug. At the end of hour 3, rats received an injection with 0.5 mg/kg of yohimbine to reinstate seeking behavior. Cue-induced seeking was evaluated during the first hour of extinction and stress-induced seeking was evaluated during reinstatement. **B. Cue-induced seeking.** Mean (± SEM) number of active and inactive spout contacts during the first hour of extinction for rats with a history of saline or heroin selfadministration pretreated with vehicle (saline) or 0.3 mg/kg LIR 6 h prior to the start of the test. **C. Extinction.** Mean (± SEM) number of active spout contacts during the three hours of extinction for the same group of rats. **D. Drug-induced seeking.** Mean (± SEM) number of active and inactive spout contacts emitted during reinstatement for the same group of rats. Saline-veh (n=6), saline-LIR (n=6), heroin-veh (n=4), heroin-LIR (n=6). *Significant differences between her-vehicle active spout contacts and all other conditions. **p<0.01; ****p<0.0001. #Significant difference within heroin-liraglutide group at different times. ##p<0.01.

### Experiment 3

#### Rotarod Test

The rotarod test was used to assess potential effects of liraglutide on locomotor performance. As evidenced in Figure 6A (left panel), liraglutide did not affect motor dexterity for a period of up to 10 hours post-injection. This was confirmed by a mixed factorial ANOVA showing no significant treatment x time interaction (F_3,35_=1.59; p>0.05), nor main effect of treatment (F<1) in the latency to fall from the rotarod at 4, 6, 8, and 10 hours after saline or liraglutide injections. As a positive control, performance on the rotarod was tested using a dose of morphine (15 mg/kg) known to reduce locomotor activity. As observed in Figure 6A (right panel), morphine clearly reduced time spent on the rotarod. This conclusion was supported by a significant main effect of treatment (F_1,10_=16.34, p<0.01.) confirming that morphine-treated rats exhibited a shorter latency to fall from the rotarod when compared with saline-treated rats overall. Further analysis was performed using survival statistics (Cox Proportional-Hazard Model logrank tests). While Figure 6B shows there were no significant differences in latency to fall between liraglutide and saline rats at 4 hours (HR_lir/sal_=1.03; CI: 0.34-3.07, p >0.05), 6 hours (HR_lir/sal_=0.8; CI: 0.28-2.3, p >0.05), 8 hours (HR_lir/sal_=1.4; CI: 0.48-4.03, p >0.05) and 10 hours (HR_lir/sal_=2.02; CI: 0.66-6.16, p >0.05), this test did show significant differences between the survival curves of rats treated with morphine and saline (Figure 6C) at 15 minutes (HR_mor/sal_=4.1; CI: 0.99-17.03, p <0.05), 30 minutes (HR_mor/sal_=4.6; CI: 01.05-20.08, p <0.05) and 60 minutes (HR_mor/sal_=3.0; CI: 0.81-11.04, p <0.05), but not at 90 minutes (HR_mor/sal_=1.78; CI: 0.55-5.79, p >0.05). Finally, Figure 6D compares survival curves during maximum effects of both liraglutide (8 hours) and morphine (15 minutes). At its greatest effect, liraglutide-treated rats fell 1/3 slower than morphine-treated rats (HR_lir/morph_=0.31; CI: 0.84-12.01, p<0.05). These results show no clear effects of liraglutide on locomotor performance.

**Figure 6.**
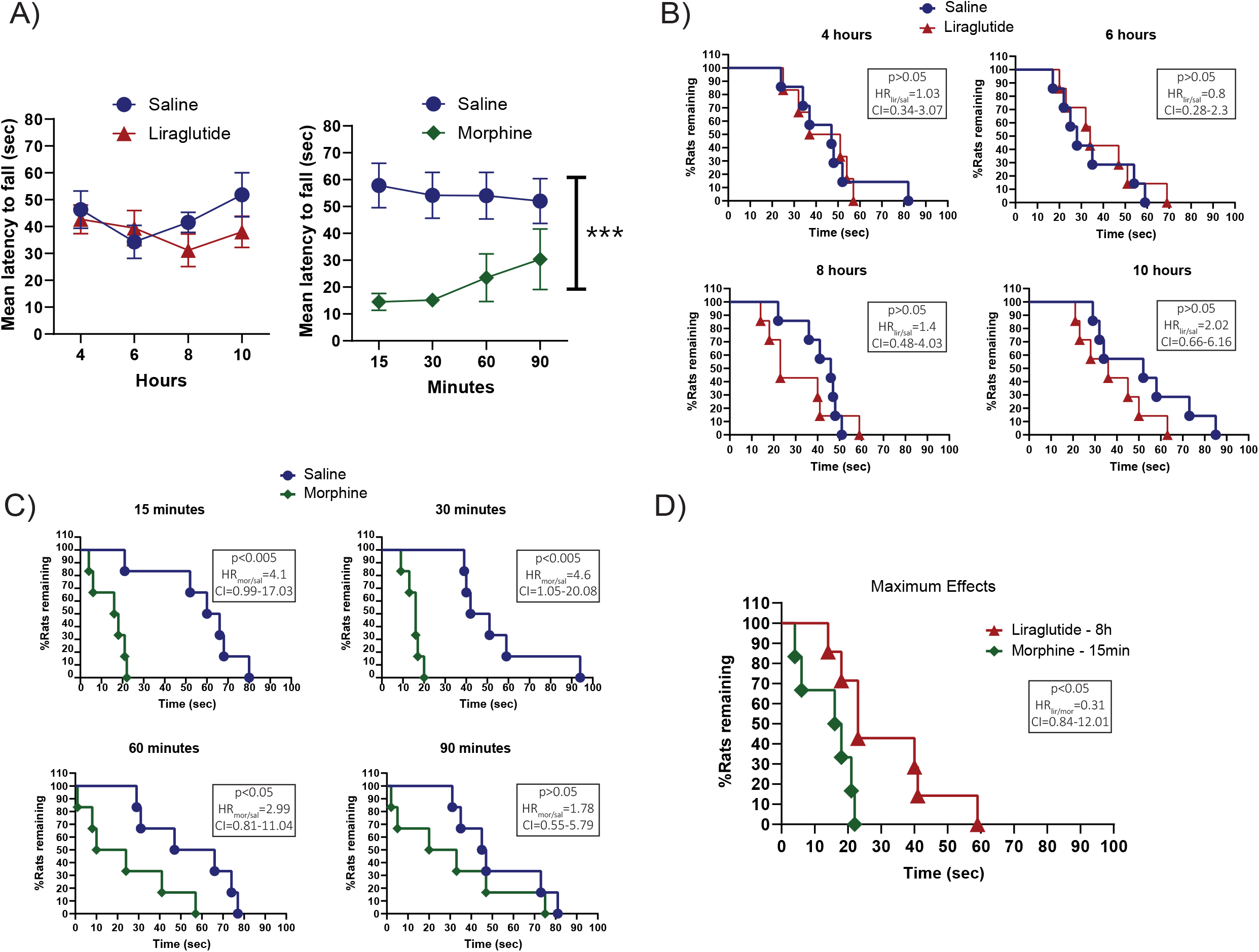
Test of motor deficiency. **A. Left Panel.** Mean (± SEM) latency to fall from the rotarod (seconds) after injection of 0.3 mg/kg liraglutide (red) or vehicle (blue) at 4, 6, 8 and 10 hours post injection. **Right Panel.** Mean (± SEM) latency to fall from the rotarod (seconds) after injection of 15 mg/kg morphine (green) or vehicle blue at 15, 30, 50 and 90 minutes post injection. **B.** Kaplan-Meier curves for latency to fall from the rotarod of rats treated with 0.3 mg/kg liraglutide (red; n=7) or vehicle (blue; n=7) at 4, 6, 8 and 10 hours post injection. **C.** Kaplan-Meier curves for latency to fall from the rotarod of rats treated with 15 mg/kg morphine (green; n=6) or vehicle (blue; n=6) at 15, 30, 60, 90 minutes post injection. **D.** Kaplan-Meier curves for latency to fall from the rotarod during maximum effects of rats treated with 0.3 mg/kg liraglutide at 8 hours post injection (red; n=7) or morphine at 15 minutes post injection (green; n=7). *Significant group effect between morphine and saline. ***p<0.001. For B, C and D the p-values indicate difference between groups in the Log-rank (Mantel-Cox) test. HR indicates the hazard ratio between the groups (indicated by subscript). CI indicates the 95% confidence interval of the ratio.

## Discussion

Research over the last decade has provided evidence that GLP-1R agonists, particularly the shorter acting Ex-4, can reduce the apparent perceived rewarding properties of drugs.^7–11,13,14,27,28^ In the present study, we showed that the acute administration of the longer-acting GLP-1R agonist, liraglutide (0.3 mg/kg, sc), can reduce cue-induced heroin seeking, drug-induced reinstatement of heroin seeking, and stress-induced reinstatement of heroin seeking behavior in rats. The effects of acute liraglutide, however, depended upon pretreatment time. Specifically, liraglutide significantly reduced cue-induced heroin seeking when evaluated six, but not four, hours post-injection. Drug-induced reinstatement was markedly reduced when assessed seven hours post liraglutide treatment; and liraglutide treatment was found to block stress-induced reinstatement of heroin seeking when administered nine hours prior to test. Taken together, these findings suggest that a minimum effective pretreatment time for liraglutide is between four and six hours in rats and that, given the appropriate pretreatment time, liraglutide can block cue-, drug-, and stress-induced reinstatement of heroin seeking in rats.

It is difficult to parse whether the temporal effects observed in these experiments are due to the liraglutide concentration achieved in blood, per se, or to the availability of the drug in brain. According to the pharmacokinetic assessment shown above, the Tmax for the 0.03 mg/kg dose in the plasma concentration curve was eight hours post-injection. However, while the 0.3 mg/kg dose of the drug was first effective behaviorally at six hours post-injection, the concentration obtained by this dose at four hours post-injection (29,618.7 ng/mL) was higher than it was at six hours (27,984 ng/mL). That said, if the AUC is proportional to bioavailability, the AUC is higher at six hours, making it possible that an AUC of at least 106,687 ngh/mL is required to reduce heroin-seeking.

As alluded to, liraglutide is a modified GLP-1 molecule that is attached to a fatty acid chain allowing it to bind to albumin to prevent degradation and release at a slower rate.^29,30^ Because the effects of GLP-1 analogs on reward-associated behaviors are centrally mediated, it is possible that, due the slower cleavage of liraglutide into its active form, the liraglutide Cmax may not correlate with the drug reaching its targets in the brain. So far, only one study has shown fluorescently labeled liraglutide (120 nmol/kg) in the hypothalamus four hours after a single iv injection.^31^ Even so, the timing remains unclear regarding the interaction of liraglutide with GLP-1Rs in the brain, particularly in nuclei associated with reward such as the nucleus accumbens and the ventral tegmental area – nuclei in which Ex-4 has proven to be effective in reducing drug-associated behaviors.^9,28^

As shown, liraglutide reduced cue-, drug-, and stress-induced heroin seeking. While it is reasonably easy to model cue- and drug-induced seeking, a number of different approaches have been used to model stress-induced seeking. The pharmacological induction of stress-like behaviors using the alpha-2 adrenergic receptor antagonist, yohimbine, has been widely described in both humans and laboratory animals;^21,22^ indicating that injection of yohimbine is a translatable method to study stress-induced reinstatement of drug-seeking behavior. Indeed, years of research have shown that yohimbine can reinstate seeking of alcohol, methamphetamine, cocaine, and nicotine in rats.^32–35^ However, some discrepancies regarding pharmacological and behavioral effects related to this method vs. natural stressors (e.g., footshock) have brought the validity of this model into question.^36^ Consequently, future studies must seek to replicate this finding using a natural stressor or injections of corticotropin releasing factor, for example^37^. Regardless, the effectiveness of liraglutide to reduce drug-seeking across all three challenges (cue, drug, and stress-induced seeking) is remarkable.

Given that the GLP-1 receptor agonist, liraglutide, reduced cue-, drug-, and stress-induced heroin seeking, consideration needed to be given to possibility that liraglutide leads to a general motor impairment or sedation. We used the rotarod to address this critical question. Importantly, the results showed that heroin-experienced rats treated with liraglutide did not differ from saline-treated controls in their performance in the rotarod test. On the other hand, rats treated with morphine—a positive control for locomotor depression—did show a shorter latency to fall compared with both saline and liraglutide-treated rats. Thus, we can conclude that the reduction in cue-, drug-, and stress-induced heroin seeking by liraglutide is not due to a simple deficit in motor coordination. A second consideration is that liraglutide may reduce responding due to gastrointestinal (GI) distress – GI distress is the most commonly reported side effects for GLP-1R agonists^38,39^. That said, and while not tested here, results from a separate set of studies showed that injection of the illness-inducing agent, LiCl, but not liraglutide (across a range of doses), increases intake of the anti-emetic clay, kaolin^18^. The longer acting GLP-1R agonist, liraglutide, then, reduces cue-induced heroin seeking, drug-induced reinstatement of heroin seeking, and stress-induced reinstatement of heroin seeking, and these effects cannot be attributed to either a generalized motor impairment or nausea.

In sum, the results found in this study are promising as they show that acute administration of the long-acting GLP-1R agonist liraglutide can not only reduce cue-induced heroin seeking, but also drug-induced reinstatement and stress-induced reinstatement of heroin-seeking in rats. It is worth noting that the different models of reinstatement are regulated by different neural mechanisms.^40^ Hence, not all compounds that reduce one are necessarily effective in reducing another. For example, current drugs used as MAT to treat OUD such as buprenorphine and methadone can reduce cue- and drug-induced seeking, but do not have an effect on stress-induced seeking.^41,42^ Orexin antagonists, on the other hand, have been shown to reduce stress and cue-induced seeking, but they are not effective in reducing drug-induced reinstatement of drug seeking behavior.^43^ Since treatments with GLP-1R agonists already are approved for the treatment of type 2 diabetes and obesity, the effectiveness of liraglutide across a wide variety of reinstatement models is a promising finding that may reveal a more effective line of treatment for OUDs in humans.^15,16^ Finally, the acute action of liraglutide provides a nonopioid option to bridge the gap for patients seeking MAT, which cannot be initiated for days following the last opioid use. Such a treatment during this period of greatest relapse risk would be of tremendous benefit to patients with OUD.

## Acknowledgements

Authors thank the National Institute on Drug Abuse for generously providing heroin. Support for this research was provided by grants from NIH DA009815 and UG3 DA050325 (to PSG) and the Pennsylvania Department of Health, Tobacco Settlement Funds SAP# 410007972 (to PSG). The authors report no financial interest or conflicts of interests.

## Authors Contribution

Respective contributions: J.E. Douton: Experimental design, data analysis and interpretation and manuscript preparation. N. Acharya: Experimental design and execution. B. Stoltzfus: experimental execution. D. Sun: HPLC/tandem MS method development and execution. P.S. Grigson: Experimental design, data interpretation and manuscript revision. JE Nyland: Experimental design, data interpretation and manuscript revision.

## Data Sharing

The data that support the findings of this study are available from the corresponding author upon reasonable request.

